# Built It, But Did They Come? Monitoring Winter Feeding Stations Use by Endangered Long-tailed Gorals

**DOI:** 10.64898/2026.01.29.702468

**Authors:** Jinhwi Kim, Donggul Woo

## Abstract

Winter supplementary feeding is widely adopted to support wild ungulates during forage scarcity, yet its ecological effectiveness for endangered mountain ungulates experiencing extreme weather–related mortality remains poorly understood. In the winter of 2023–2024, a mass mortality event of long-tailed gorals (*Naemorhedus caudatus*) in South Korea prompted the installation of emergency feeding stations in core habitats. We used camera traps to quantify feeding activity and assess how environmental and demographic factors shaped visitation. Long-tailed gorals fed primarily around midnight, deviating from their typical diurnal and crepuscular patterns. Feeding frequency increased when dried mulberry (*Morus alba*) leaves was available, snow covered the ground, and fawns were present. Conversely, visitation declined with warmer temperatures and rainfall. Agonistic interactions peaked during periods of feed depletion, indicating heightened competition over limited resources. Although several individuals occasionally gathered, most stations consistently supported only one or two individuals, reflecting limited reach at the population level. This study provides the first systematic assessment of endangered Long-tailed gorals’ behavioral and temporal responses to winter supplementary feeding, offering critical baseline data to evaluate and optimize conservation interventions for mountain ungulates.

---

Supplementary feeding is a widely used wildlife management strategy intended to provide nutrition when natural forage is scarce (Inslerman and Baker 2006). Its primary goals include improving animal health, enhancing reproductive output, and supporting population stability (Peterson and Messmer 2007, Robb et al. 2008, Newey et al. 2010). Wildlife managers have applied this practice to both game and non-game species to mitigate the effects of habitat degradation (Miranda et al. 2015, Borowski et al. 2019), seasonal food shortages (Page and Underwood 2006), and other environmental stressors (Bramorska et al. 2023, Tóth et al. 2023). Particularly in temperate regions, deep snow and prolonged cold restrict forage access (Warchałowski et al. 2015, Jackson et al. 2021), prompting programs designed to reduce overwinter mortality and support reproduction (Inslerman and Baker 2006, Bishop et al. 2009, Horstkotte et al. 2020). For example, *ad libitum* feeding substantially improved survival and body condition in mule deer (*Odocoileus hemionus*) and pronghorn (*Antilocapra americana*) during severe winters (Peterson and Messmer 2007, Bishop et al. 2009).

Despite its short-term benefits, supplementary feeding entails ecological risks. First, uptake is inconsistent; some animals never visit feeding sites, and even those that do may consume little of the provided food (Katona et al. 2014). Consequently, in some cases, supplementary feeding yields no observable improvement in body condition or reproductive output (Putman and Staines 2004). Even when animals do use feeders, repeated access can alter movement patterns and foraging behavior (Peterson and Messmer 2007, Bramorska et al. 2023, Saldo et al. 2024). Furthermore, feeding sites may become hotspots for disease transmission (Cotterill et al. 2018)—as observed in the spread of chronic wasting disease among cervids (Huang et al. 2025)—and can contribute to localized overgrazing (Van Beest et al. 2010). Finally, the long-term conservation outcomes of supplementary feeding remain poorly understood (Newey et al. 2010, Kubasiewicz et al. 2016).

The long-tailed goral (*Naemorhedus caudatus*) is an endangered mountain ungulate vulnerable to habitat loss and winter mortality (Roy et al. 1995, OH et al. 2025). In South Korea, extreme winter conditions have occasionally caused catastrophic die-offs. Historical records document over 6,000 deaths in 1967 (Won 1967). More recently, in early 2024, a sudden freeze following rainfall led to the death of an estimated 1,042 individuals—nearly half the national population (OH et al. 2025). To mitigate further loss, the Ministry of Climate, Energy and Environment installed emergency feeding stations in key goral habitats.

We used camera traps to investigate the feeding and behavioral dynamics of long-tailed gorals at these emergency feeding stations. Specifically, we aimed to (1) characterize temporal patterns of visitation; (2) quantify the influence of environmental drivers (snow, temperature) and resource availability on feeding intensity; (3) document the presence of non-target species to assess potential disease risks; and (4) examine agonistic interactions to evaluate intraspecific competition. Our findings aim to establish an evidence-based foundation for optimizing supplementary feeding practices for endangered mountain ungulates.

## STUDY AREA

We conducted monitoring in the Samcheok and Uljin regions of South Korea (**Figure 1**), where supplementary feeding for long-tailed gorals was initiated following the discovery of >30 carcasses during the winter of 2010 (Park and Hong 2021). Although past estimates suggest approximately 100 individuals inhabited the area in the early 2000s (Yang 2002), winter mortality remains a recurring threat. Most recently, 68–74 goral deaths were reported during the 2023–2024 winter (Green Korea United 2024), prompting the Ministry of Climate, Energy and Environment to implement emergency feeding measures. Five new feeding stations were installed across the two regions, and provisioning began on 11 December 2024. The average elevation of the feeding sites was 780 m in Samcheok and 471 m in Uljin.

**Figure 1.**
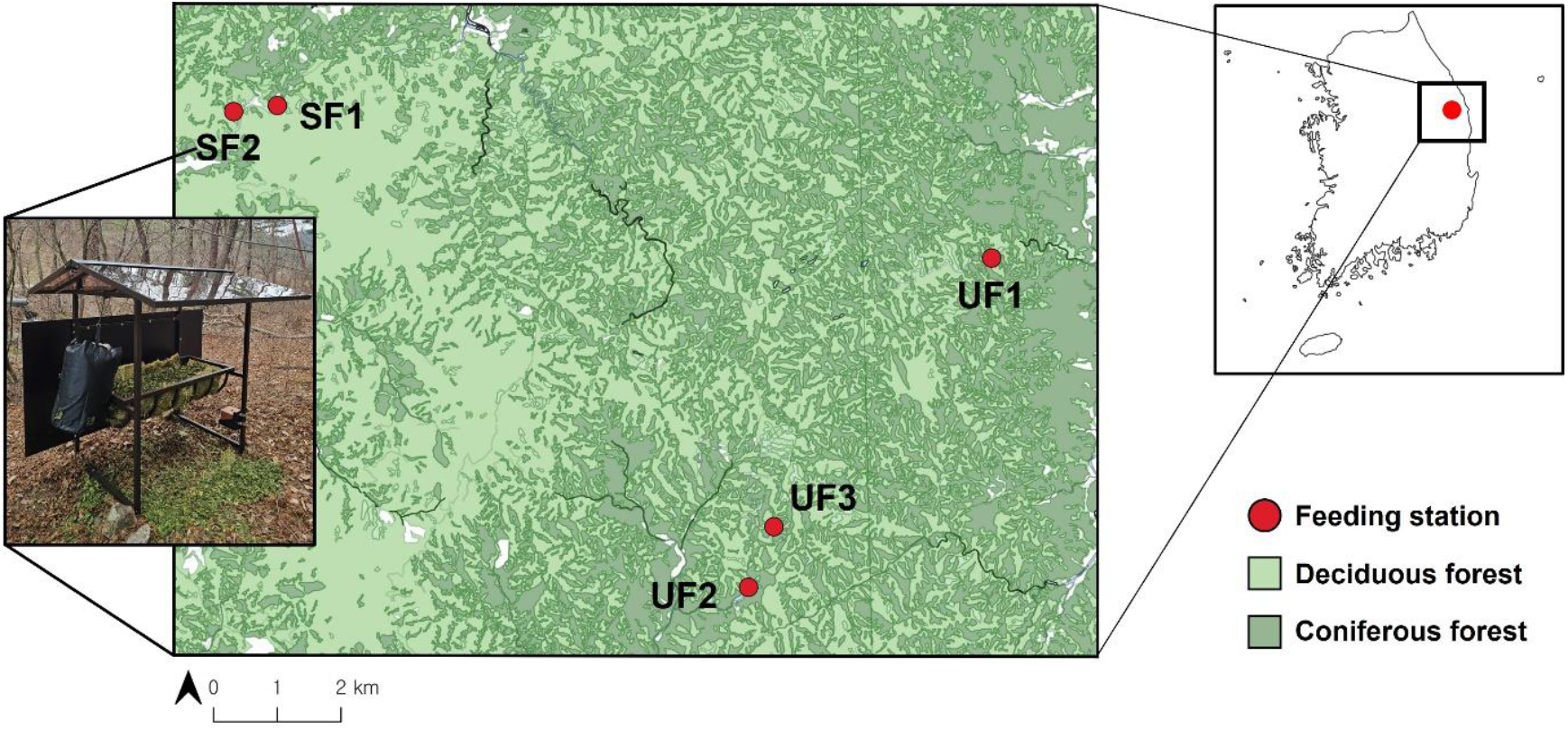
Map of the study area (Samcheok and Uljin) in Republic of Korea. The map shows five newly installed long-tailed goral feeding stations (red circles) monitored during the 2024– 2025 winter, overlaid on the distribution of deciduous and coniferous forest types. The photograph shows the feeding station setup monitored in this study

We obtained weather data from the Korea Meteorological Administration via the nearest Automatic Weather Stations: Seokpo (37.04°N, 129.00°E) for Samcheok and Geumgangsong (36.94°N, 129.25°E) for Uljin. Both stations are located approximately 14 km from the respective feeding sites. During the study period (December 2024–April 2025), Samcheok recorded a mean monthly temperature of -0.2°C (range -6.1–9.9°C) and a mean monthly precipitation of 24.5 mm (range 0–73.5 mm). In Uljin, the average monthly temperature was 3.9°C (range -0.5–12.6°C) with mean monthly precipitation of 34.3 mm (range 0–107.5 mm).

## METHODS

### Camera trap installation and monitoring

To monitor goral activity, we deployed trail cameras (Reconyx Hyperfire2; Reconyx, Inc., Holmen, WI, USA) at each station starting 11 December 2024. We equipped each station with 2–3 cameras positioned 4–5 m from the feeder to capture entering and exiting animals. Cameras recorded either 3 consecutive images (1-s interval) or 30-s dynamic video clips triggered by motion. We visited stations biweekly to replace batteries and memory cards, and to replenish feed (15 kg dried mulberry [*Morus alba*] leaves) and mineral blocks (sodium chloride, calcium, vitamin A). We recorded the time to first detection to assess the initial latency of station use.

### Data processing

We extracted date and time from image metadata. Individual identification was not feasible due to the lack of distinct natural markings or horn features visible in nocturnal footage. Therefore, we analyzed data at the event level. We defined a feeding event as a series of detections separated by <5 minutes. Following Popova et al. (2017), we truncated events at 30 minutes.

For each detection, we identified the species and recorded the maximum number of individuals observed simultaneously. We coded mulberry provision, snow cover, and fawn presence (juveniles <1 year) as binary variables (1= present, 0 = absent). We classified events as feeding if animals consumed mulberry or licked the ground or mineral blocks. Agonistic behaviors (threat, fight, defense) were classified following Voloshina and Myslenkov (2012).

### Statistical analysis

We paired the 281 feeding events recorded between December 2024 and April 2025 with daily average temperature and precipitation data. To estimate the effects of environmental and management factors on daily feeding activity, we used a Bayesian negative binomial regression model implemented in the brms package (Bürkner 2017) in R version 4.3.0 (R Core Team, 2023).

We modeled expected daily feeding events using a log link function. Predictors included standardized temperature and precipitation, snow cover, fawn presence, and mulberry provision, plus station and month. We assumed equal detectability across stations and included no offset terms. We present results from the additive model, as interaction terms did not improve model fit based on leave-one-out cross-validation (LOO-CV). We applied weakly informative priors: Normal(0, 1.5) for intercept and fixed effects, and Gamma(0.01, 0.01) for the shape parameter. Posterior samples were drawn using Markov Chain Monte Carlo (MCMC) via Stan (four chains of 4,000 iterations each, 2,000 warm-up). All parameters showed convergence (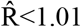, effective sample sizes > 1,000). We report posterior medians and 95% credible intervals (CrIs, 2.5th–97.5th percentiles) to summarize effect sizes and uncertainty.

To estimate the minimum number of individuals using feeding stations per day, we modeled the daily maximum count (n = 289) using a Bayesian Poisson mixed-effects model with station as a random intercept. We used weakly informative priors (Normal(0, 1.5) for the intercept; half-Student-t(3, 0, 10) for group-level standard deviation) and the same MCMC settings. Station-level estimates were summarized by posterior medians and 95% CrIs.

## RESULTS

### Feeding site use

Long-tailed gorals first visited the feeding stations between 4–39 days after installation. Subsequently, they fed regularly on the day of, or the following, replenishment. On average, feeding activity was recorded on 56.2 days per station (range: 38–77 days). Snow was present on an average of 19.6 of those days (34.9%), and precipitation occurred on 6.4 days (11.4%). We recorded 2,182 goral detections, of which 1,760 (80.7%) involved feeding (**Figure 2(A)**). Feeding activity peaked around midnight (23:00–01:00) and gradually declined to a minimum around midday (10:00–14:00). During peak hours, Samcheok stations recorded approximately twice as many feeding events as Uljin stations, though both regions exhibited similar diel pattern with reduced visitation around midday. Weekly trends (**Figure 2(B)**) indicated that Samcheok stations were visited more frequently in January and February. In contrast, Uljin stations (UF1, UF2, UF3) exhibited delayed but more pronounced utilization, with feeding activity increasing markedly in early March and peaking around Week 11 (March 10–16; weekly total: 156 events across three stations). Subsequently, visitation declined sharply across all sites during late March and early April, coinciding with rising temperatures (mean daily temperature >10°C), snowmelt, and the emergence of natural spring vegetation.

**Figure 2.**
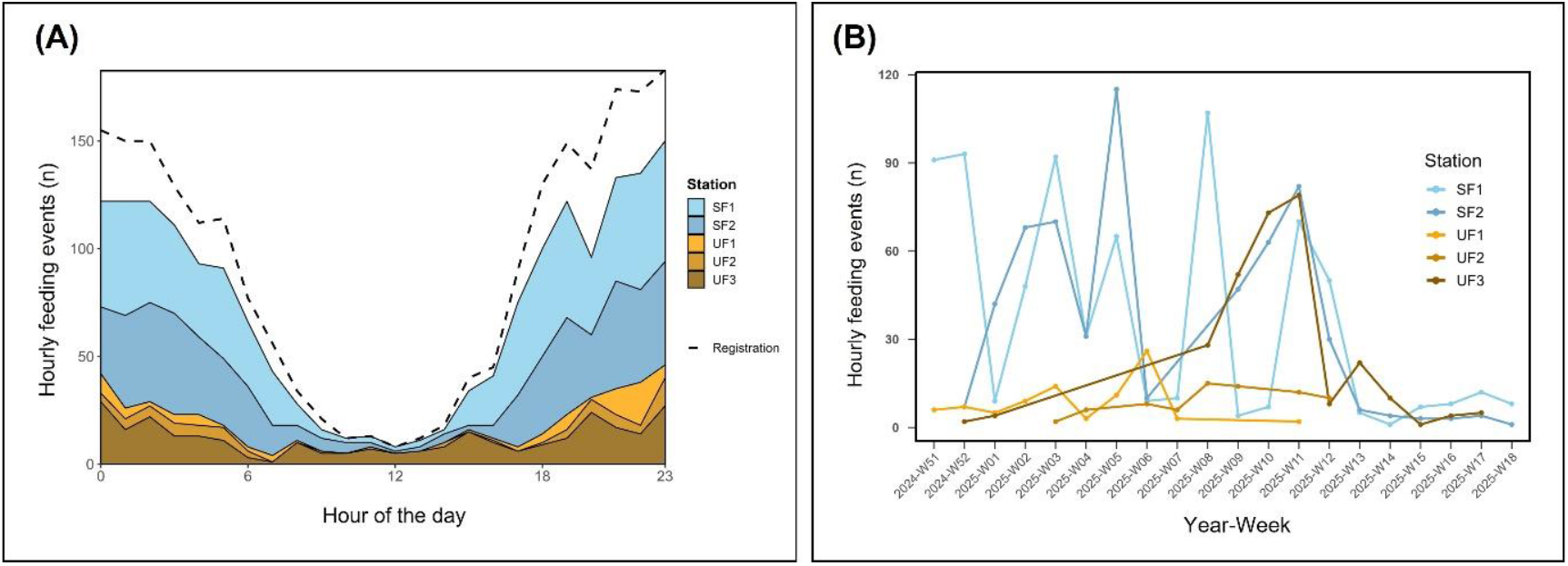
Temporal patterns of Long-tailed goral feeding events and registrations at feeding stations during winter 2024–2025. (A) Stacked area plot showing the distribution of hourly feeding events (n = 1,770) at stations in Samcheok (blue) and Uljin (yellow) regions. The dashed line presents total event registration (n= 2,182), including feeding, passing, and roaming. (B) Weekly trends in hourly feeding events by stations. Lines present the number of feeding events recorded per week at each station, highlighting differences in feeding intensity and timing across regions and stations.

We recorded 76 detections of non-target wildlife. Visits were rare from December to March (1– 5 detections per month), but increased markedly in April (n = 36), primarily involving Asian badgers (*Meles leucurus*) (Appendix A). We recorded 3 wild boar (*Sus scrofa coreanus*) detections but no visits from sympatric water deer (*Hydropotes inermis*) or roe deer (*Capreolus pygargus tianschanicus*) at any station.

### Effect of environmental and management factors

We modeled daily feeding activity using a Bayesian negative binomial regression (**Figure 3, Figure 4**). Among all predictors, mulberry provision had the strongest positive effect on feeding activity (β = +1.3; 95% CrI: 1.0–1.5), followed by fawn presence (β = +0.8; 95% CrI: 0.6–1.0) and snow cover (β = +0.2; 95% CrI: 0–0.4). Conversely, average temperature (β = -0.02; 95% CrI: -0.05–0.00) and precipitation (β = -0.03; 95% CrI: -0.05–0.00) showed weak but consistent negative associations. Station and month effects exhibited broader credible intervals, reflecting greater uncertainty.

**Figure 3.**
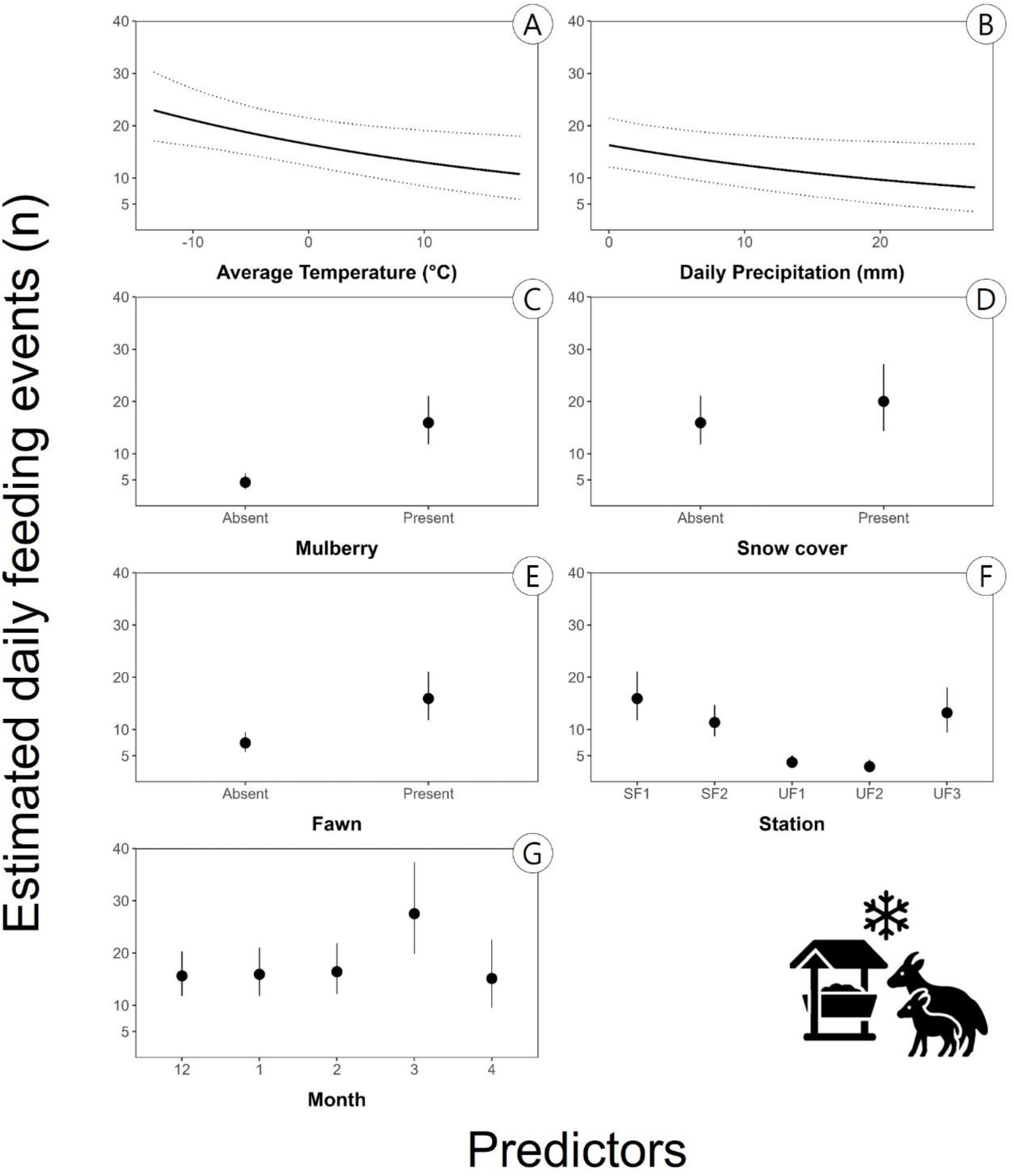
Effects of environmental and management factors on daily feeding events of long-tailed goral at supplementary feeding stations from December 2024 to April 2025. Estimates derive from a Bayesian negative-binomial regression (n= 281 feeding events). The plots show predicted daily feeding events in relation to predictors (A–G). In panels A and B, solid lines denote posterior medians and dotted lines 95% credible intervals; in panels C–G, points and vertical bars show posterior medians and 95% credible intervals.

**Figure 4.**
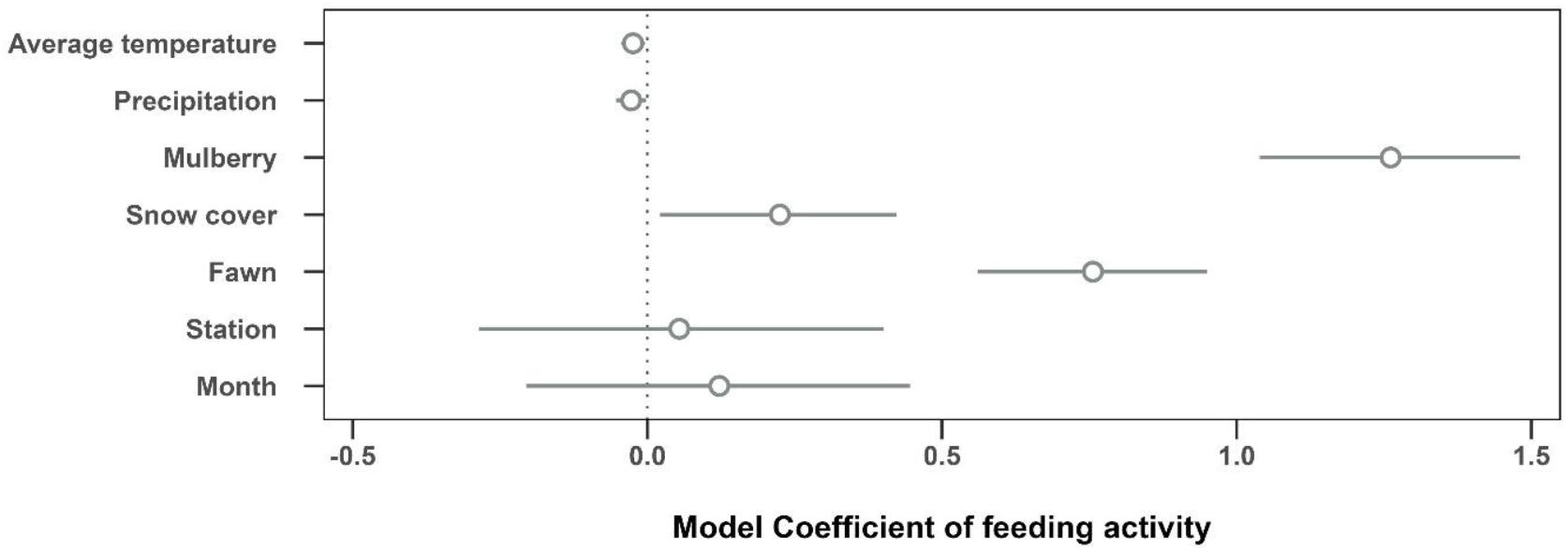
Estimated effects of environmental and management factors on the daily number of feeding events by long-tailed goral. Points represent the median posterior estimates, and horizontal lines indicate the 95% credible intervals from the Bayesian negative binomial regression model. Predictors include average temperature, daily precipitation, mulberry provision, snow cover, fawn presence, station, and month.

### Minimum daily visitors from daily maximum counts

Daily maximum counts of gorals varied across stations (**Figure 5**). For clarity, SF and UF refer to feeding stations located in Samcheok and Uljin, respectively. Observed mean daily maximum counts (range) were: SF1, 2.3 individuals (1–7); SF2, 1.9 (1–6); UF1, 1.6 (1–5); UF2, 1.1 (1–3); and UF3, 1.3 (1–4). Posterior median estimates from the Bayesian Poisson model aligned with these observations: SF1, 2.3 individuals (95% credible interval [CrI]: 1.4–3.8); SF2, 1.9 (95% CrI: 1.1–3.0); UF1, 1.6 (95% CrI: 1.0–2.6); UF2, 1.1 (95% CrI: 0.7–1.6); UF3, 1.3 (95% CrI: 0.8–2.1). The model estimated substantial overlap in station-level variability (σ_Station = 0.5, 95% CrI: 0.2–1.2).

**Figure 5.**
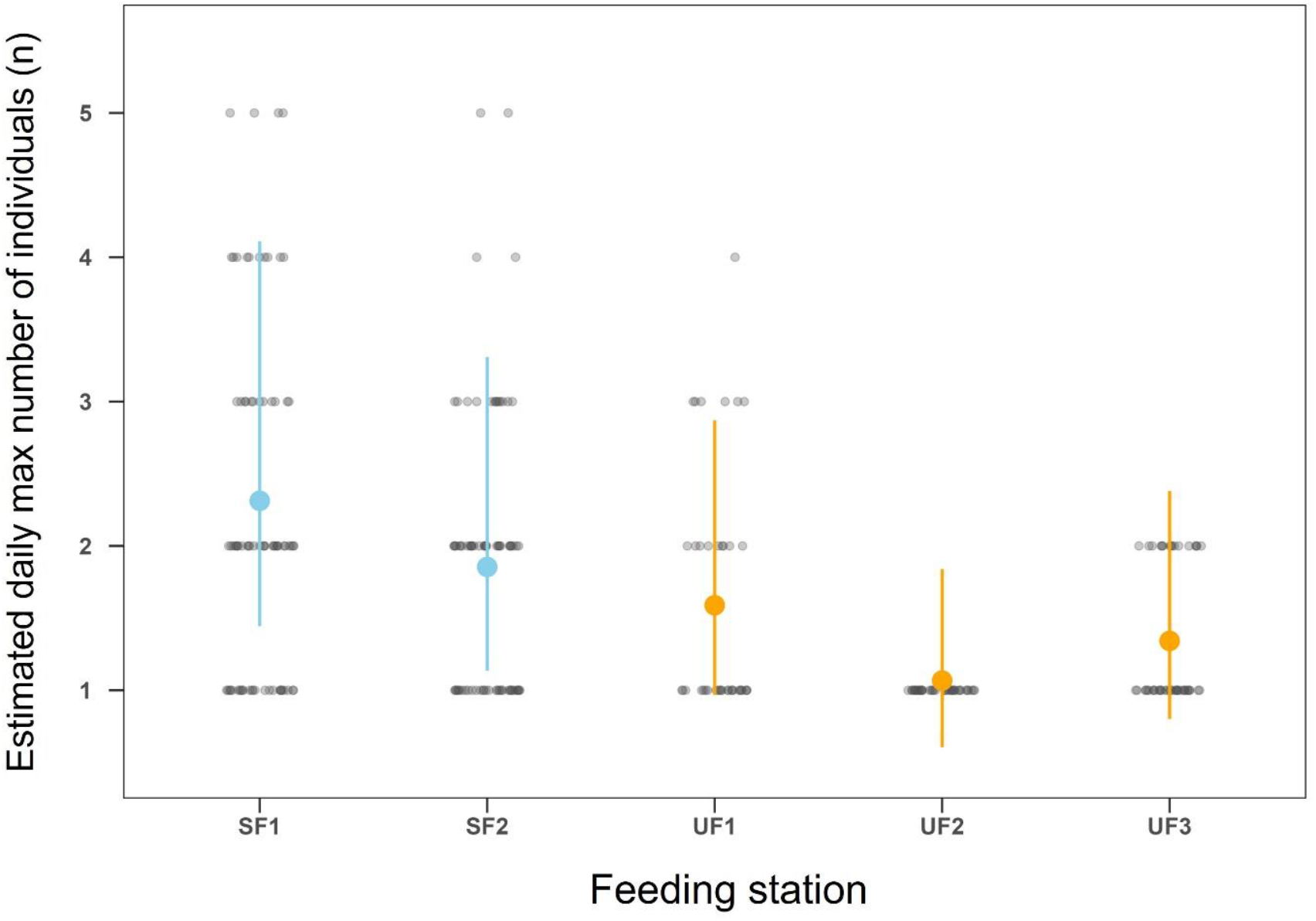
Posterior estimates of the estimated daily maximum counts by station. Colored points (blue for Samcheok stations; yellow for Uljin stations) and vertical lines show the posterior medians and 95% credible intervals for the expected daily maximum counts of long-tailed gorals at each feeding station. Overlaid grey dots show the observed daily maximum counts.

### Agonistic behavior

We recorded 851 agonistic behaviors: 363 threats, 293 fights, and 195 defense events **(Figure 6)**. We observed no agonistic interactions after Week 14 (early April), which coincided with the sharp decline in goral visitation. Daily counts peaked on 24 December 2024 (102 events), 15 January 2025 (96), and 16 January 2025 (58). These peaks coincided with feed depletion events (indicated by vertical dashed lines in Figure. 6), suggesting a link between resource limitation and intraspecific conflict.

**Figure 6.**
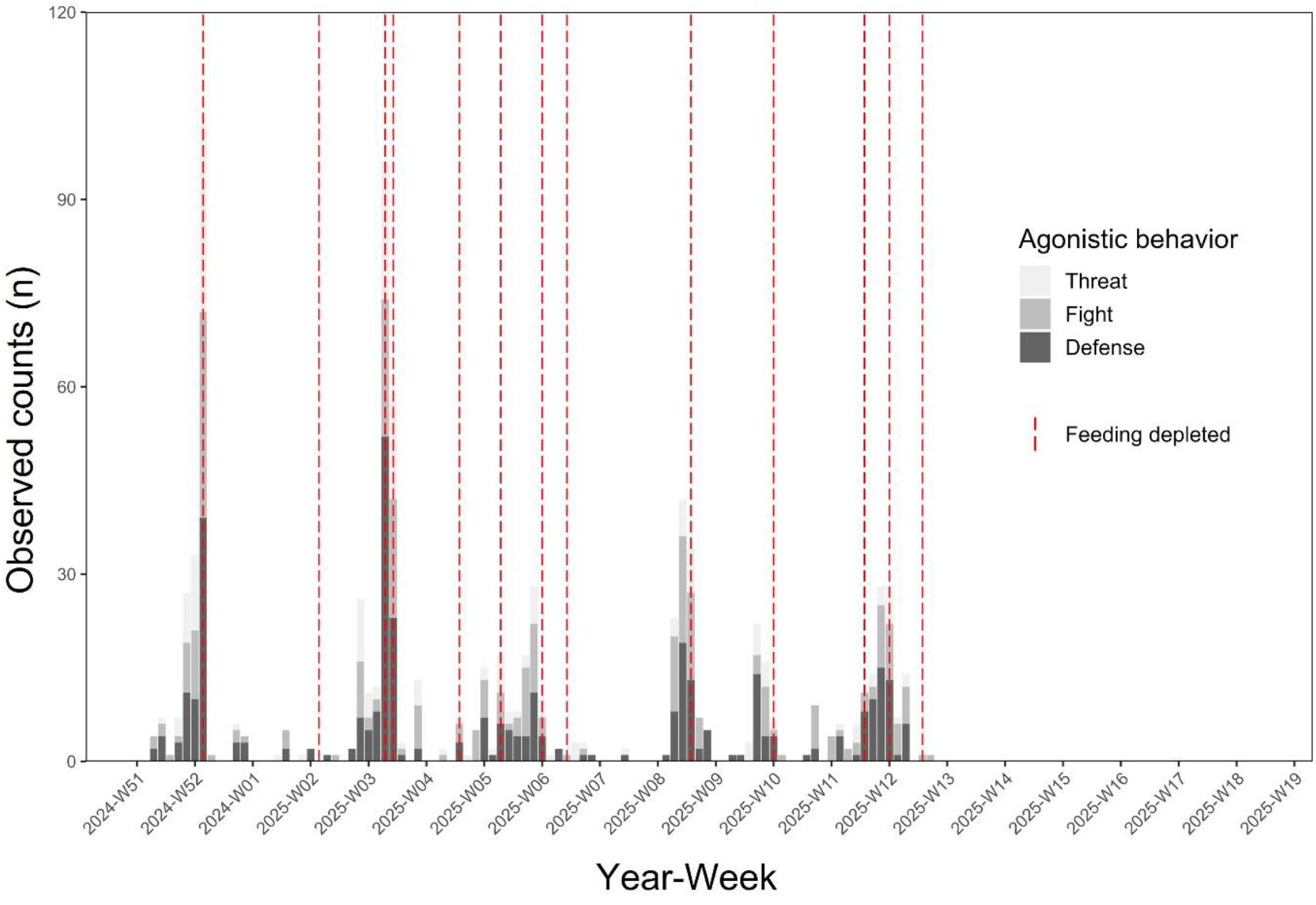
Daily counts of agonistic behaviors (Threat, Fight, Defense) observed at feeding stations from December 2024 to April 2025 (n = 851). Red dashed lines indicate dates when feed was depleted at one or more stations. Bar heights reflect summed data across all stations.

## DISCUSSION

Although winter supplementary feeding has been implemented in South Korea for nearly a decade, its ecological effectiveness for endangered long-tailed gorals has remained largely unassessed. This study provides the first empirical evaluation of these emergency interventions. Our monitoring demonstrates that while gorals use supplementary feed, particularly during deep snow and periods of high energetic demand, the conservation benefits are modulated by distinct behavioral shifts, intraspecific competition, and limited reach at the population level. Our monitoring revealed that goral visitation to supplementary feeding stations peaked around midnight and declined sharply after dawn. This pronounced nocturnal activity contrasts with the typically crepuscular or diurnal patterns reported for long-tailed gorals (Lim et al. 2025) and the bimodal activity of Himalayan gorals (Anwar 1989). While Voloshina and Myslenkov (2012) observed active midday foraging in winter, the absence of such behavior in our study suggests that gorals were either utilizing natural vegetation elsewhere or resting to minimize energy expenditure. This shift to nocturnal station use likely reflects an adaptive strategy to mitigate winter energy deficits. As ungulates downregulate their metabolic rates (Timmons et al. 2010, Arnold 2020) and reduce movement to conserve energy (Shakeri et al. 2021, Ahn 2022, Bramorska et al. 2023), gorals may prioritize predictable, high-value food during the coldest hours. Specifically, this behavior may optimize thermoregulation via the heat increment of feeding. Since ruminant digestion generates peak metabolic heat approximately 4–6 hours post-ingestion (Smith et al. 1978), midnight feeding strategically aligns this internal heat production with the lowest ambient temperatures (03:00–07:00). Similar to observations in moose (*Alces alces*) and red deer (*Cervus elaphus*) (Arnold et al. 2004, Rea et al. 2013), our findings suggest that goral nocturnal activity is a physiological response to thermal stress rather than a primary tactic for predator avoidance.

Agonistic interactions were most frequent during late December and mid-January, coinciding with periods of feed depletion. Although most feeding stations were typically used by only one or two individuals, these peaks in conflict also coincided with times when the highest numbers of individuals were observed. This suggests that both resource scarcity and temporary aggregation increased the likelihood of direct encounters. Similar patterns have been observed in chamois (*Rupicapra rupicapra*) and llama (*Lama glama*), where aggressive interactions increased under conditions of high resource quality and availability, consistent with a resource-defense strategy (Fattorini et al. 2018, Panebianco et al. 2021). However, the severity of these interactions was generally mild. Long-term observations on goral populations by Voloshina and Myslenkov (2012) reported only a few cases of injurious conflict in this species, suggesting that most agonistic behaviors function as low-level competitive displays rather than escalated aggression. Nevertheless, subordinate individuals may still experience elevated stress or exclusion, underscoring the importance of maintaining a consistent feed supply to reduce conflict and ensure equitable access.

Long-tailed gorals visited feeding stations more frequently on colder days with little precipitation, though these effects were modest. In contrast, snow cover had a stronger positive effect, likely due to limited natural forage availability. Similar patterns have been reported in other ungulates: roe deer increased feeder use under deep snow (Ossi et al. 2017, Tóth et al. 2023), and European bison (*Bison bonasus*) remained closer to feeders during snowy periods (Bramorska et al. 2023). Snow not only reduces forage access but also increases locomotion costs, further discouraging natural foraging (Shakeri et al. 2021). In rugged terrain, such stress may have severe demographic consequences, as seen in adult chamois, whose mortality increases under prolonged snow cover (Rughetti et al. 2011). For long-tailed gorals, snow likely represents a key ecological threshold that shifts the cost-benefit balance in favor of artificial provisioning.

Fawn presence was also positively associated with station use, suggesting increased activity by adult females with offspring or possibly pregnant females. Adult females with offspring are constrained by the energetic demands of gestation and the limited mobility of juveniles, making them more likely to exhibit fidelity to predictable food resources. Carcass records from Uljin highlight this vulnerability: 89.4% of gorals found dead in 2010 were yearlings or pregnant females (Park and Hong 2021). Supplemental feeding can help mitigate this risk; previous studies have shown that enhanced winter nutrition substantially improves juvenile survival (Bishop et al. 2009), and supports neonatal viability for pregnant females (Jackson et al. 2021). These findings underscore the importance of maintaining winter food availability for vulnerable reproductive cohorts.

Visits by non-target wildlife were limited, particularly during early winter. Most occurrences were recorded between March and April, coinciding with the decline in goral visitation. Notably, sympatric ungulates such as roe deer and water deer were not observed, and wild boar detections were limited to brief passing events after March. This suggests the current feeding protocol was selective. The provision of dried mulberry leaves—a forage less attractive to generalists than grains or root vegetables—likely minimized multi-species aggregation (Campbell et al. 2013, Popova et al. 2017). Additionally, the relatively high elevation of feeding sites likely restricted access by competitors until temperatures rose. Continued monitoring is needed to confirm the long-term consistency of this species-specificity.

Although some stations occasionally hosted larger groups, most were consistently used by only one or two individuals. This suggests that the current program may have limited population-level reach, potentially due to monopolization by dominant individuals or groups. Similar limitations have been noted in other ungulate and alpine herbivore studies, where supplementary feeding benefited specific individuals without yielding broader demographic responses (Newey et al. 2010). Addressing such spatial and social inequalities is essential to maximize conservation value of winter provisioning. This study provides the first systematic evaluation of winter feeding for long-tailed gorals in South Korea. However, as our findings rely on a single season, multi-year monitoring is necessary to confirm the consistency of these responses. Future efforts should also incorporate dietary analysis (e.g., fecal sampling) to quantify the actual nutritional contribution of supplementary feed relative to natural forage.

## MANAGEMENT IMPLICATIONS

Our results demonstrate that long-tailed gorals do use supplementary feeding stations— particularly during periods of snow cover, when natural forage availability is reduced. Agonistic behaviors intensified around times of feed depletion, highlighting the potential for conflict when provisioning is inconsistent. These findings suggest that ensuring a reliable feed supply during severe winter conditions may reduce intraspecific competition and support the survival of vulnerable individuals. To maximize effectiveness, feeding programs should be designed to provide timely and consistent access during critical periods (Inslerman and Baker 2006, Ewen et al. 2015), especially when snow depth impedes natural foraging.

Regarding specific protocols, we recommend continuing the use of dried mulberry leaves, as this diet effectively attracted gorals while minimizing non-target visits and potential disease transmission. Furthermore, because stations were often dominated by a small number of individuals, increasing the number of feeding sites rather than the volume of food at single sites, may better serve the wider population. While supplementary feeding can be a valuable short-term strategy, it should be implemented as part of a broader management approach that considers habitat quality, landscape connectivity, and the species’ long-term ecological needs.

## ACKNOWLEDGMENTS

We thank the Korea National Park Service for their essential support in installing the supplementary feeding stations, and the Ministry of Climate, Energy and Environment for granting permission to conduct this research. We are also grateful to the staff of the Wangpicheon Environmental Station for their assistance in replenishing feed supplies at key sites. Finally, we acknowledge the Mammal Restoration Team at the National Institute of Ecology for their dedicated help with feeding station maintenance, including the replacement of SD cards and consistent field support throughout the monitoring period. This study was funded by the Ministry of Climate, Energy and Environment (NIE-C-2025-78).

## ETHICS STATEMENT

This study was conducted using non-invasive monitoring methods, including motion-activated camera traps positioned at a distance from feeding stations to avoid disturbing the animals. No animals were captured, handled, or otherwise manipulated during the research period. The placement and operation of camera traps were designed to minimize human interference, ensuring that the behavior and feeding patterns of long-tailed gorals were observed under natural conditions.

## DATA AVAILABILITY STATEMENT

The data and R code that support the findings of this study are openly available in the Open Science Framework (OSF) at https://doi.org/10.17605/OSF.IO/BN2XT.

---

**APPENDIX A..**
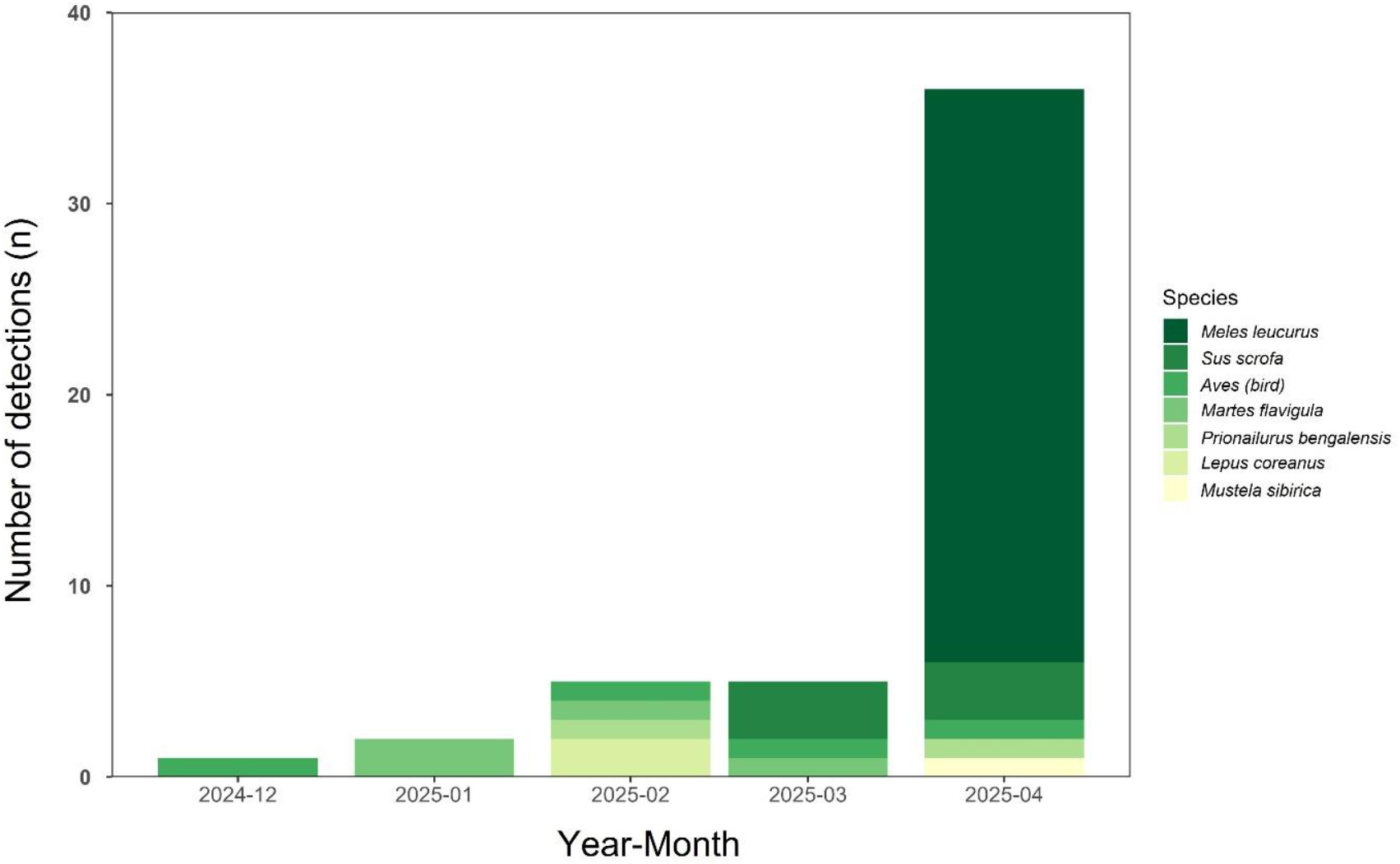
Monthly detections of non-targeted species at feeding stations (December 2024– April 2025). The stacked bar plots show the number of detections of non-targeted species per Year-Month (n = 76).

## Notes

https://osf.io/bn2xt/

### Competing Interest Statement

The authors have declared no competing interest.

